# Resistance to the conjugation inhibitor AZT reshapes metabolic compatibility between mating partners during plasmid transfer

**DOI:** 10.64898/2026.07.13.738253

**Authors:** Eda Deniz Erdem, Alan X. Elena, Peiju Fang, Thomas U. Berendonk, Uli Klümper

## Abstract

Plasmid-mediated dissemination of antibiotic resistance genes (ARGs) is a major driver of antimicrobial resistance (AMR). The nucleoside analogue zidovudine (azidothymidine, AZT) has emerged as a promising conjugation inhibitor, yet the consequences of AZT resistance for subsequent plasmid transfer remain poorly understood. Here, we investigated how AZT resistance in donor and recipient bacteria influences conjugative plasmid transfer. Independent AZT-resistant mutants arose through nonsense mutations in two distinct thymidine metabolism genes, *yjjG* and *tdk*. Consistent with previous studies, AZT suppressed plasmid transfer 10-fold between susceptible mating partners, whereas resistance in both partners restored transfer despite continued AZT exposure. Unexpectedly, plasmid transfer was markedly reduced even in the absence of AZT when only one mating partner carried a resistance mutation, declining by up to 11.5-fold regardless of whether the resistant strain was the donor or the recipient. This mixed-resistance phenotype was rescued by supplementation with dTMP, a thymidine pathway metabolite downstream of both resistance-associated mutations, thereby increasing plasmid transfer by up to 5.88-fold and restoring transfer to levels comparable to those of susceptible-susceptible matings. In contrast, excess dTMP reduced conjugation between susceptible partners, supporting a broader role for balanced nucleotide metabolism in determining conjugation efficiency. Together, our findings identify nucleotide metabolism as a previously unrecognised determinant of conjugative plasmid transfer and suggest that the long-term efficacy of conjugation inhibitors such as AZT will depend not only on the evolution of resistance, but also on how resistance reshapes the physiological compatibility of bacterial mating partners.

**Importance:** Antibiotic resistance spreads rapidly because conjugative plasmids transfer resistance genes between bacteria. Drugs that inhibit plasmid transfer are therefore being explored as a new strategy to limit antimicrobial resistance, but little is known about how resistance against these compounds affects subsequent gene transfer. We show that resistance to the conjugation inhibitor zidovudine (AZT) does not simply restore plasmid transfer. Instead, transfer depends on the resistance status of both mating partners. While two susceptible or two resistant bacteria transferred plasmids efficiently, mixed susceptible-resistant pairings showed markedly reduced conjugation even in the absence of AZT. Restoring thymidine metabolism rescued this phenotype, identifying balanced nucleotide metabolism as a previously unrecognised determinant of conjugative plasmid transfer. These findings suggest that the long-term performance of conjugation inhibitors will depend not only on the evolution of resistance, but also on how resistance reshapes the physiological compatibility of bacterial mating partners, providing a new framework for understanding and designing anti-conjugation strategies.

## Main text

The global rise of bacterial antimicrobial resistance (AMR) is partly driven by horizontal transfer of antibiotic resistance genes (ARGs) (1,2). Conjugation, the direct transfer of mobile genetic elements (MGEs) between bacterial cells, majorly contributes to this process, enabling ARG dissemination across species boundaries (3,4). As a substantial proportion of ARGs are encoded on transferable plasmids (5), targeting conjugation represents a promising but underexplored strategy to limit AMR spread (6,7).

The thymidine nucleoside analogue zidovudine (3′-azido-3′-deoxythymidine, AZT) was the first antiretroviral drug approved for HIV/AIDS treatment in 1987 (8). Following cellular uptake, AZT is activated through the thymidine salvage pathway by thymidine kinase (TdK), enabling its incorporation into DNA, where it acts as a chain terminator (9,10). Recently, AZT has attracted attention for its antibacterial activity, particularly against members of the Enterobacteriaceae (8,9), and, at subinhibitory concentrations, a candidate conjugation inhibitor, that suppresses plasmid transmission (11–13). Nonetheless, bacteria readily acquire AZT resistance, most commonly through spontaneous mutations in the *tdk* gene. This prevents AZT activation through phosphorylation and can abolish its antibacterial and conjugation-inhibiting activity (8,11). Restoration of plasmid transfer has been demonstrated when both mating partners gain AZT resistance (11,13), while the other consequences of resistance, for example in only one partner, remain unknown.

To understand how resistance evolution influences the long-term efficacy of conjugation inhibitors, we investigated how AZT resistance in donor and recipient cells affects subsequent conjugative plasmid transfer in presence and absence of AZT (Supplementary Methods). We used a chromosomally *mCherry*-tagged *Escherichia coli* donor carrying the broad-host-range IncP-1 tetracycline-resistance plasmid pKJK5::*gfp* (3) and a non-fluorescent, gentamicin-resistant *E. coli* recipient (14), allowing donor, recipient, and transconjugant cells to be distinguished by selective plating and fluorescence (15). Spontaneous AZT-resistant donor and recipient mutants were generated by incubation on LB agar supplemented with AZT (32 µg/mL; 4×MIC), with gained resistance confirmed by MIC determination (TableS1). Genome comparisons between AZT-resistant strains and their susceptible counterparts identified resistance-associated SNPs in two independent genes linked to thymidine metabolism (TableS2). In the resistant recipient, a nonsense mutation in *tdk* introduced a premature stop codon that truncates TdK (Fig.1A), likely rendering it non-functional through loss of its active site and disruption of proper folding (16,17). In the resistant donor a stop-gain mutation occurred in *yjjG*, which encodes a deoxyuridine monophosphate (dUMP) phosphatase that is part of the *de novo* dTMP generation pathway, while also hydrolysing non-canonical pyrimidine derivatives as part of its house-cleaning function (Fig.1A) (18–20). The precise functional consequence of this mutation, aside from gained resistance, remains unclear, but altered YjjG activity could potentially affect nucleotide salvage, dUMP accumulation, or AZT-derived substrates. Importantly, no resistance-associated fitness loss was observed in the absence of AZT for either mutation with growth kinetics not substantially altered (Fig.S1).

**Figure 1:**
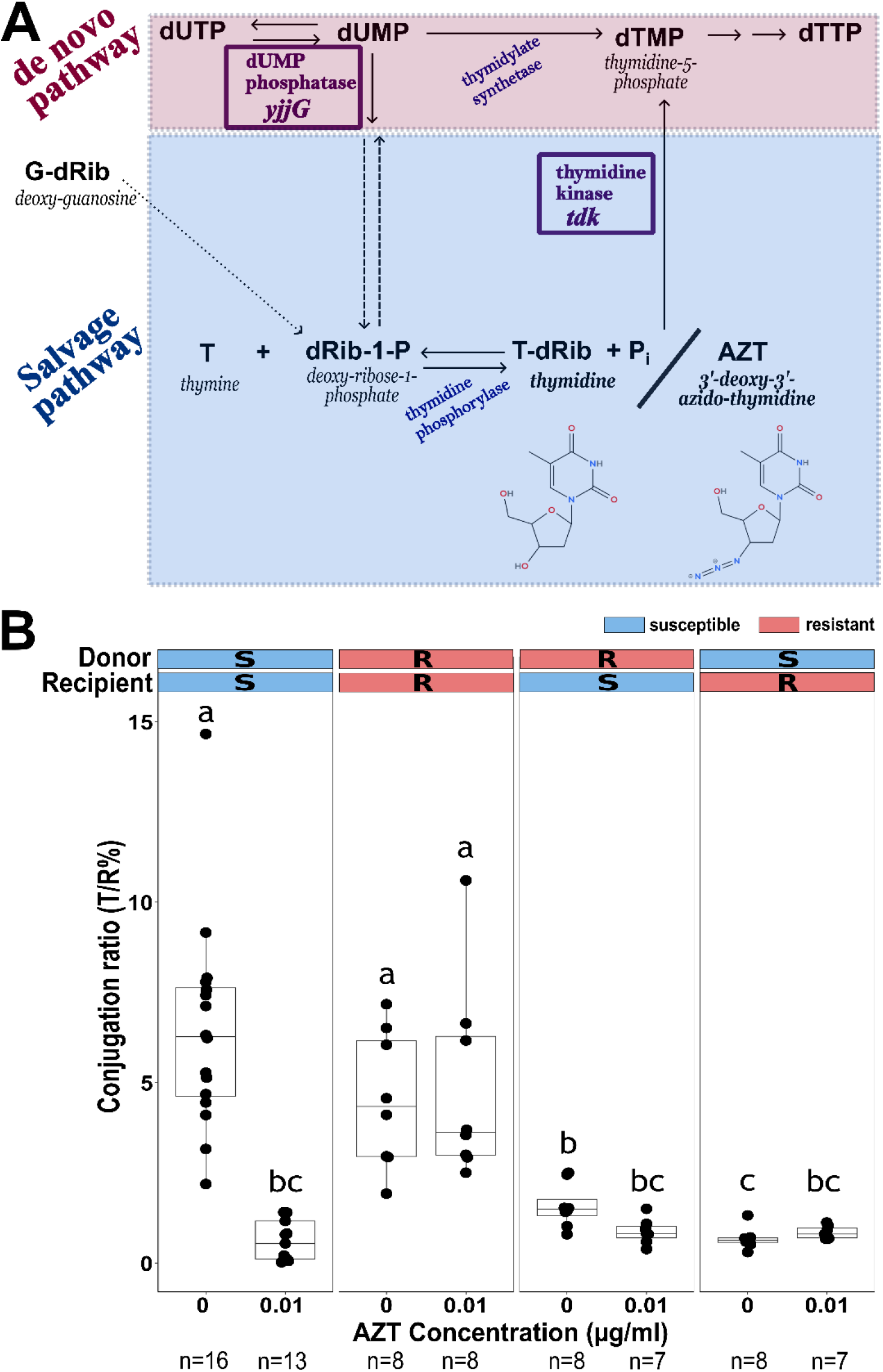
Plasmid transfer rates in presence and absence of AZT. (A) Schematic of the thymidine salvage (blue) and de novo synthesis (pink) pathways in *E. coli*, adapted from Itsko & Schaaper, 2011 (20). Purple boxes highlight *tdk*, encoding thymidine kinase, and *yjjG*, encoding dUMP phosphatase, the two genes identified by genome comparison to carry resistance-associated SNPs in the AZT-resistant recipient and donor strains, respectively. TdK activity is required for the phosphorylation of AZT to its active form, making it essential to AZT’s mechanism of action. Loss or impairment of YjjG activity may disrupt the salvage pathway or cause dUMP accumulation, potentially overactivating thymidylate synthase and rendering the salvage pathway dispensable. (B) Conjugation efficiency across mating conditions, attributable to AZT exposure as well as resistance profiles of the mating partners (as indicated on top of the graph), after a liquid mating period of 24 h. The conjugation ratio was calculated as the normalised ratio of transconjugants per recipient, after selective plating. The number of biological replicates per condition is shown below each group. Differing letters denote statistically distinct groups following Kruskal–Wallis rank-sum test with Dunn’s post hoc pairwise comparisons and Benjamini–Hochberg correction for multiple testing.

We quantified plasmid transfer among all susceptible and resistant donor-recipient combinations in the presence and absence of AZT using semi-growth-dependent mating assays (Supplementary Methods), with transfer frequency expressed as transconjugants per recipient. Plasmid transfer differed strongly across mating conditions (χ^2^=59.75, *P*<0.0001, Kruskal–Wallis rank-sum test; Fig.1B, Supplementary Data). In susceptible donor-recipient matings, transfer occurred at 6.44±2.81% transconjugants per recipient in the absence of AZT but declined 10.4-fold to 0.62±0.54% under AZT exposure (FDR-adjusted *P*<0.00001). This reduction was not accompanied by changes in plasmid copy number per cell (Fig.S2; *P*=0.09, Mann-Whitney U test), suggesting that AZT suppresses conjugation through mechanisms other than impaired intracellular plasmid replication.

AZT resistance in both mating partners abolished AZT-mediated suppression of conjugation, with transfer frequencies remaining similar in AZT absence and presence (4.53±1.78% vs. 4.88±2.59%; FDR-adjusted *P*=0.49), comparable to those of susceptible mating partners in AZT absence (*P*=0.22 & 0.24; Fig.1B). Similarly, bacterial densities displayed no AZT-dependent reduction in resistant donor-recipient matings (Supplementary Data; *P*>0.1). Thus, our system reproduced the established pattern that AZT suppresses transfer between susceptible mating partners, whereas resistance in both partners restores transfer under AZT exposure (11,12). Despite arising through distinct genetic routes, the two resistance mutations were functionally compatible.

Unexpectedly, even in the absence of AZT, this compatibility was lost when only one mating partner carried a resistance mutation, regardless of the mutated gene (Fig. 1B, Supplementary Data). Relative to susceptible-susceptible matings, plasmid transfer without AZT dropped 4.1-fold with only a resistant donor (1.58±0.56%, FDR-adjusted *P*<0.01) and 9.6-fold with only a resistant recipient (0.67±0.27%, *P*<0.0001). AZT addition did not further affect plasmid transfer in either of these mixed-resistance pairings, with transfer frequencies remaining similarly low as for susceptible-susceptible matings in AZT presence (*P*>0.05). Conjugation frequency was nonetheless slightly but significantly higher when only the donor, rather than only the recipient, carried the resistance mutation (*P*=0.048), suggesting that the mixed-pair transfer frequency may be affected by both the distinct thymidine-metabolism functions of the mutated genes (16–19) and the different roles of donor and recipient cells during plasmid transfer (21).

The reduced conjugation frequency in AZT absence observed between mixed-resistance mating partners suggests that resistance-associated perturbations in thymidine metabolism may underlie the observed transfer phenotype. To test this hypothesis, we examined plasmid transfer across strains with different resistance profiles under dTMP supplementation. As a metabolite downstream in the pyrimidine biosynthesis pathway, dTMP bypasses the metabolic steps affected by both *tdk* and *yjjG* mutations (22). dTMP supplementation significantly influenced plasmid transfer (χ^2^=48.35, *P*<0.0001; Kruskal-Wallis rank-sum test; Fig.2A, Supplementary Data). Reduced conjugation frequency between mixed-resistance mating partners was rescued regardless of whether the resistant partner was the donor or the recipient, increasing transfer by 2.43- and 5.88-fold, respectively (FDR-adjusted *P=*0.04 & 0.004). Indeed, plasmid receipt increased to levels comparable with susceptible-susceptible baseline matings (*P*>0.05).

**Figure 2:**
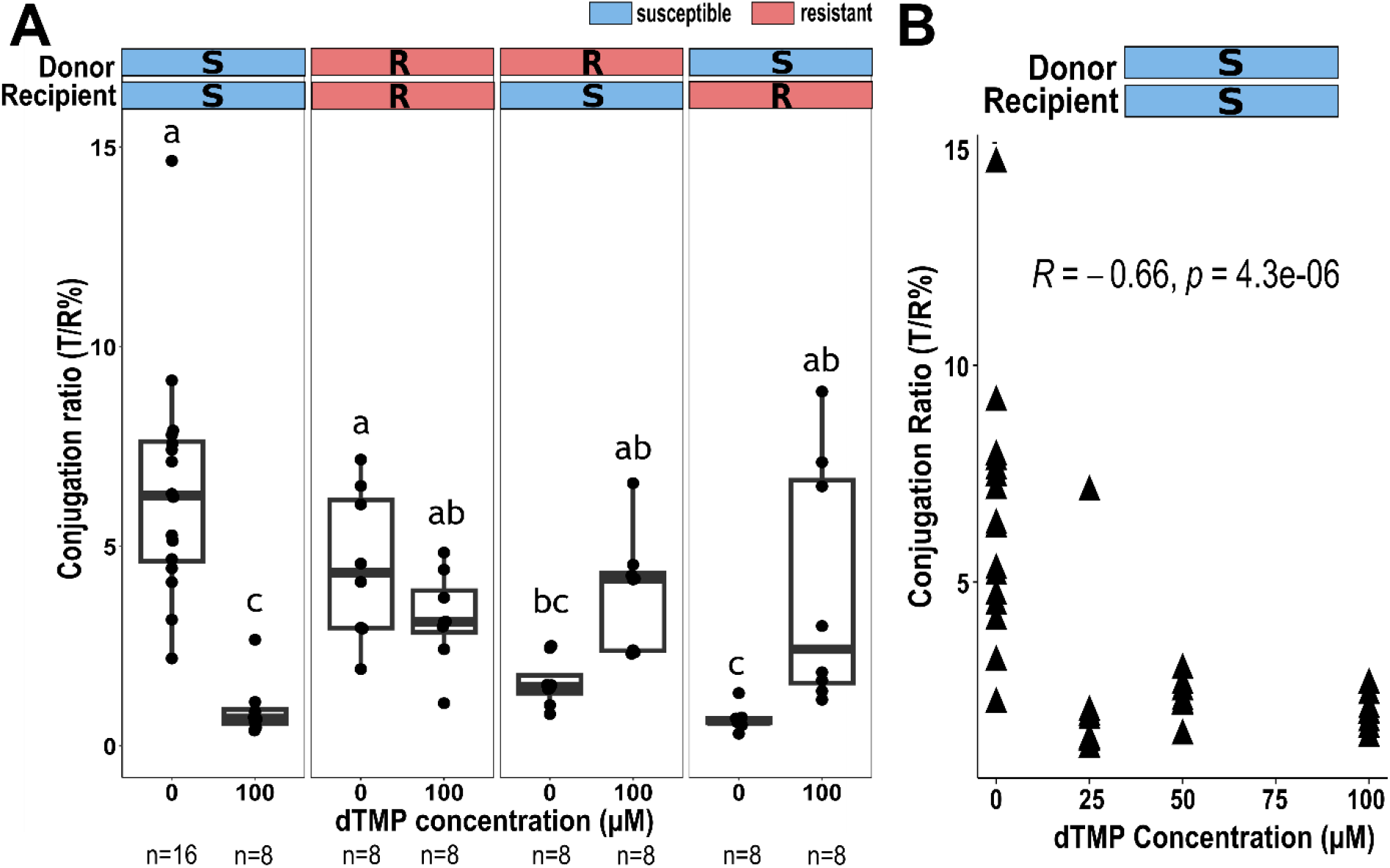
Plasmid transfer rates in presence and absence of dTMP supplementation. Conjugation efficiency across mating conditions, attributable to dTMP exposure as well as resistance profiles of the mating partners (as indicated on top of the graph) (A) The effect of dTMP on conjugation efficiency for all mating pairs at 100 µM. The number of biological replicates per condition is shown below each group. Letters denote statistically distinct groups following Kruskal–Wallis rank-sum test with Dunn’s post hoc pairwise comparisons and Benjamini–Hochberg correction for multiple testing. (B) Spearman correlation analysis between dTMP concentration and the rate of plasmid transfer efficiency between susceptible-susceptible mating pairs.

Conversely, dTMP did not significantly affect plasmid transfer or final transconjugant density when both mating partners were resistant (*P=*0.27 & 0.34), but decreased conjugation efficiency by up to 8.34-fold in susceptible donor-recipient matings from 6.44±2.81 to 0.92±0.69% in a concentration-dependent manner (Rs=-0.66, *P*<0.0001, Spearman rank correlation; Fig.2B). This reduction may result from disturbances in intracellular deoxynucleotide pools caused by excess thymidine, generating nucleotide imbalance that can fundamentally alter cellular physiology (23). Consistent with this concept, perturbations in cellular metabolism have increasingly been recognised as important determinants of conjugative plasmid transfer and its associated physiological costs (24,25).

Our findings indicate that AZT-induced suppression of plasmid transfer requires susceptibility in both mating partners and that resistance-associated mutations in two distinct genes involved in thymidine metabolism can generate a donor-recipient incompatibility phenotype that compromises conjugation efficiency. Circumventing the metabolic consequences of these mutations by dTMP supplementation restored plasmid transfer, supporting nucleotide metabolism as the primary mediator of the observed phenotype. These findings further suggest that targeting a single step in thymidine metabolism is unlikely to provide an evolutionarily robust anti-conjugation strategy, as resistance can rapidly restore transfer. Instead, future anti-conjugation approaches may benefit from simultaneously targeting multiple metabolic steps within the thymidine pathway to increase the evolutionary barrier to resistance. Together, our findings identify balanced nucleotide metabolism as a previously underrecognized determinant of conjugative plasmid transfer and suggest that the long-term efficacy of anti-conjugation strategies will depend not only on the evolution of resistance itself, but also on how resistance reshapes the physiological compatibility of bacterial mating partners. Rather than identifying AZT as a standalone anti-conjugation therapy, this work highlights nucleotide metabolism as a promising target space for the development of next-generation conjugation inhibitors that may prove more robust to resistance evolution.

## Acknowledgements

The authors thank Christiane Zschornack, Melanie Tannert, and Steffen Kunze for technical support in the laboratory. This work was supported by the JPIAMR TEXAS, the JPIAMR SEARCHER, project here funded by the German Bundesministerium für Forschung, Technologie & Raumfahrt under grant numbers 01KI2401 & 01KI2404A and the Urban Resistome project funded by the Deutsche Forschungsgemeinschaft (DFG) under project number 460816351. UK was supported through the ONE-BRIDGE project funded by the EU4Health Program (EU4H) under project number 101233407. UK & EDE received support through the DAAD-funded JELLY-AMR project (Project ID: 57747282). AXE and TUB were supported through the RHUMARGE project funded by the Deutsche Forschungsgemenischaft (Grant number 544004729). PF was supported through the China Scholarship Council (CSC) under grant number 202004910327. Responsibility for the information and views expressed in the manuscript lies entirely with the authors.

## Data Availability

The datasets supporting the conclusions of this article are included within the article and its additional files or available through the corresponding author upon reasonable request. Original sequencing data is available in the NCBI sequencing read archive under project accession number PRJNA1492545.

## Author Contributions

**Eda Deniz Erdem:** Conceptualization; Data curation; Formal analysis; Investigation; Methodology; Validation; Visualization; Writing – original draft; Writing – review & editing

**Alan X. Elena:** Data curation; Formal analysis; Funding acquisition; Investigation; Project administration; Software; Writing – review & editing

**Peiju Fang:** Investigation; Methodology; Writing – review & editing

**Thomas U. Berendonk:** Conceptualization; Funding acquisition; Project administration; Resources; Supervision; Writing – review & editing

**Uli Klümper:** Conceptualization; Data curation; Funding acquisition; Methodology;

Project administration; Resources; Supervision; Visualization; Writing – review & editing

