## Supplementary information for "Resistance to the conjugation inhibitor AZT reshapes metabolic compatibility between mating partners during plasmid transfer"

### Supplementary Methods

#### 1. Bacterial strains

The antibacterial and conjugation-inhibiting effects of AZT on Enterobacterales are well established, particularly against *E. coli*, the focal strain of this study (1–5). The donor strain chosen for the plasmid transfer experiments, *E. coli* MG1655::*mCherry* carrying the plasmid pKJK5::gfpmut3-35, is chromosomally tagged with a gene cassette, $lacI_q$-pLpp-mCherry-$Km^R$, which provides kanamycin (Km) resistance and red fluorescence. For the recipient strain, a wild-type *E. coli* MG1655 strain chromosomally tagged with the gentamicin (Gm) resistance gene *aacC1* was used (6). The plasmid pKJK5::gfpmut3-35 encodes tetracycline (Tet) resistance and green fluorescent protein (GFP), which can only be expressed when the plasmid is transferred to the recipient since the aforementioned chromosomal $lacI_q$ repressor is absent (7, 8). This allowed for the differentiation of donor, recipient, and transconjugant bacteria by selective plating and distinct fluorescence characteristics. MICs of AZT for susceptible donor and recipient strains were determined by the broth dilution method as described by (9).

Resistance to AZT in donor and recipient strains was generated by incubating susceptible bacteria on LB agar plates containing 32 µg/ml AZT (4× MIC; Thermo Scientific, Waltham, MA, USA) for 48 h at 37°C, as this approach has been reported to successfully induce mutations in the *tdk* gene (1, 2). Among the colonies that emerged, a single colony was selected and cultivated for each of the donor and recipient strains. AZT resistance of the resulting strains was confirmed by determining MICs using the broth microdilution method, in accordance with EUCAST guidelines (10).

#### 2. Bacterial Growth Dynamics under AZT Presence

Bacterial growth kinetics analyses were performed to determine whether AZT affected the growth kinetics of susceptible and resistant strains at low concentrations, thereby establishing suitable AZT concentrations for the plasmid transfer experiments. These analyses were also used to examine the potential fitness costs that AZT resistance may impose in the absence of AZT in resistant strains, compared with susceptible strains.

Overnight liquid cultures of all bacterial strains were washed, resuspended, and diluted with PBS to a bacterial concentration of $10^8$ colony-forming units (CFU)/mL. 6 µl of bacterial suspensions were added to 294 µl of sterile LB medium containing 0.0, 0.001, 0.1 and 0.1 µg/ml of AZT in a sterile 96-well plate, along with negative controls only containing liquid media. For each condition, 3 technical replicates were fashioned. The plate was incubated for 24 h at 37°C with continuous shaking in a microplate reader, and optical density (OD) at 600 nm was measured every 15 minutes. These measurements were used to build a logistic model and calculate growth dynamics parameters, such as the area under the curve (AUC) and fastest generation time (doubling time, $T_{gen}$).

##### 3. DNA extraction

Total bacterial DNA for each of the 4 strains was extracted from 2 mL of liquid cultures after washing and resuspending the bacteria in PBS, using MasterPure Complete DNA and RNA Purification Kit (Lucigen, Middleton, WI, USA) according to the manufacturer's protocol. The isolated DNA was assessed for quantity and quality using a NanoDrop™ (Thermo Scientific) and stored at -20°C for further analyses.

##### 4. Whole Genome Sequencing

To identify resistance-conferring mutations in AZT-resistant strains and determine whether genes other than the previously reported *tdk* were affected, strains were whole-genome sequenced. Genomic DNA libraries were constructed with the Illumina DNA Library Prep Tagmentation low-volume protocol and sequenced on an Illumina NovaSeq 6000 employing NovaSeq S4 v1.5 4XP chemistry (200-cycle kit) to produce paired-end 2 × 100 bp reads. Demultiplexing was executed via bclconvert version 4.4.6. Sequencing produced approximately 4.7-7.4 million read pairs per isolate.

Samples were sequenced using a NovaSeq 6000 and a 2x150 bp approach. Raw reads were trimmed using Trimmomatic V0.39 (11) using a quality threshold of 30. Quality-optimised reads were used to evaluate the presence of mutations in resistant donor and recipient strains. Briefly, the closest available high-quality reference genome was found by calculating the hash distance between test strains and a set of sketched RefSeq genomes (12, RefSeq-release 229). The selected RefSeq strain was used as the reference genome for SNP calling using Snippy (13). After this, variable regions resulting from transposition events were identified and removed using Gubbins (14).

The biological impact of the detected SNPs was assessed with SnpEff (15). Synonymous mutations were not taken into consideration for further downstream analysis. To properly identify mutations rising from AZT exposure, resistant strains were compared to their susceptible counterparts in pairs. Only those mutations present in the resistant strain but not in the susceptible strain were analysed in depth.

#### 5. Conjugative mating assay

To examine the role of AZT as a conjugation inhibitor and the impact of resistance on its efficacy, a semi-growth-dependent conjugative mating assay, based on our existing growth-dependent mating protocol (11, 12), was developed by increasing the initial bacterial concentration in the mating systems. This modification was made to ensure that the AZT-resistant partner would not outcompete the AZT-susceptible partner in the presence of AZT (0.01 µg/ml) for mating pairs with mixed resistance profiles. This strategy, as an alternative to growth-independent methods, considered the active cellular machinery and alterations in the bacterial metabolic state when evaluating the rate of plasmid transfer, while evading the growth-limiting effects of AZT. For plasmid transfer experiments using deoxythymidine monophosphate (dTMP, Thermo Scientific), the same protocol was followed.

Prior to the mating assay, bacterial liquid cultures were incubated at 37°C, shaking at 140 rpm overnight in LB medium containing Km at 50 µg/ml and Tet at 10 µg/ml for the donor strains, and Gm at 20 µg/ml for the recipient strains. The strains were washed and resuspended in PBS, then diluted to an OD600 of 1. The assays were conducted with 7 to 16 biological replicates in glass vials containing 20 ml of sterile LB medium, to which donor and recipient strain solutions were added at a 1:1 ratio. The final concentrations in the mating systems were $5x10^7$ CFU/ml per strain. The mating mixtures were incubated at 37°C for 24 h, shaking at 140 rpm, and after serial dilution with PBS, inoculated on LB agar plates with corresponding antibiotics (Km 50 µg/ml and Tet 10 µg/ml for selection of donor bacteria, Gm 20 µg/ml for selection of recipient bacteria, and Gm 20 µg/ml and Tet 10 µg/ml for selection of transconjugant bacteria). The conjugation ratio was calculated by normalising the final conjugation of transconjugants to that of recipients.

**6. Assessment of plasmid replication rate via quantitative PCR**

To determine if the AZT's effect on plasmid transfer is associated with alterations to the plasmid replication machinery, possible changes to the relative plasmid abundance in *E. coli* monoculture in the presence of AZT were examined. For this analysis, *E. coli* strain CM2372, a lacZY deletion mutant of *E. coli* MG1665 with IncP1-α plasmid pG527 encoding kanamycin resistance, was used. Specific primers and probe sequences designed to target the pG527 plasmid as an amplicon are available at (16).

Six replicates of liquid cultures of *E. coli* CM2372 were incubated at 37°C, shaken at 140 rpm with and without AZT exposure (0.01 µg/mL). Upon reaching the maximum carrying capacity, DNA was extracted as described above. The plasmid prevalence was determined by normalising pG527 abundance to 16S rRNA gene abundance, with all quantifications performed via real-time qPCR on a C1000 Touch™ Thermal Cycler (Bio-Rad, Hercules, CA, USA). For the pG527 plasmid amplicon target, reactions were set up by adding template DNA (10 ng/µL), primer pairs (300 nM), and the detection probe (80 nM) to 10 µl of Luna® Universal Probe qPCR Master Mix (New England Biolabs, Massachusetts, USA). For the 16S rRNA gene, 10 µL of Luna® Universal qPCR Master Mix (New England Biolabs) and 300 nM of the primer pair 338F (CCTACGGGAGGCAGCAG) - 518R (ATTACCGCGGCTGCTGG) (17) were used, and the probe was omitted. Each reaction was carried out in technical triplicates with a final volume of 20 µL. The cycling conditions consisted of an initial denaturation step for 10 min at 95°C, followed by 44 cycles of 15 s at 95°C and 30 s at 60°C. The recombinant plasmid pBELX-1 extracted from *E. coli* CM2455 (18) was used in concentrations of $10^8$-$10^2$ copies per reaction to build the standard curve. Accepted results were those with standard curve amplification efficiency between 0.9 and 1.1, $R^2$ values equal to or greater than 0.99, and no multiple peaks in the melting curve analysis. A tenfold dilution of template DNA demonstrated no PCR inhibition.

**7. Statistical analysis and Data Visualisations**

All statistical analyses, data manipulation, and visualisation were performed in RStudio v2026.01 (19, 20). Multiple comparisons were conducted using the Kruskal–Wallis rank-sum test followed by Dunn's test for post hoc analysis (21) and pairwise comparisons were carried out using the Mann–Whitney U test. Multiple testing correction was applied using the Benjamini–Hochberg false discovery rate method via the multtest R package (22), with significance defined as $P \leq 0.05$.

Correlations were evaluated using Spearman's rank method and visualised with ggpubr (23). The R package growthcurver was used to plot growth curves and estimate growth dynamics parameters (24), and all further visualisations were produced using ggplot2 (25).

### Supplementary Figures

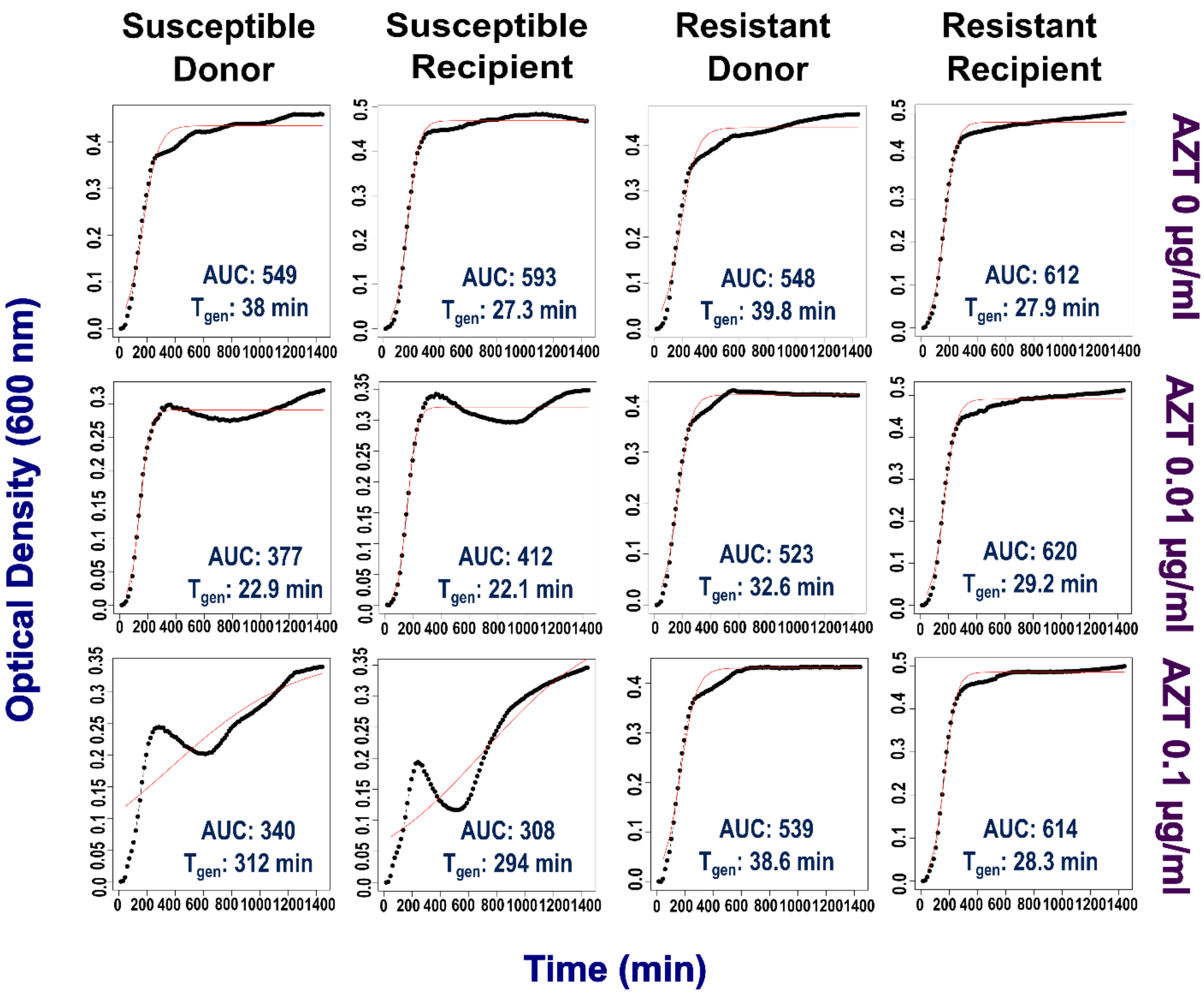


**Fig S1: Growth curves of bacterial strains in the presence of AZT** at concentrations of 0.0, 0.01, and 0.1 µg/ml for 24 hours at 37 °C, represented as optical density ($OD_{600}$) over time (min). Black lines indicate experimental results averaged from three technical replicates; red lines illustrate logistic model fits estimated by growthcurver (24), which was utilised to derive the minimum generation time ($T_{gen}$) and area under the curve (AUC) parameters of growth kinetics.

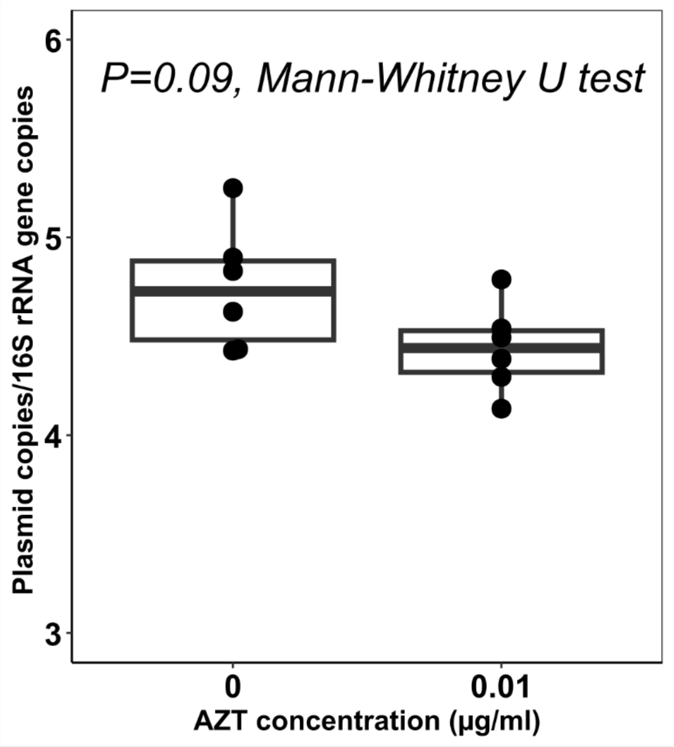


**Fig S2: Plasmid copy number normalised to 16S rRNA gene copy number in *E. coli* CM2372**, a lacZY deletion mutant of *E. coli* MG1655 carrying the IncP-1α plasmid pG527, following incubation with AZT (0.01 μg/mL) and without AZT presence to the maximum carrying capacity. AZT exposure led no significant effect on plasmid copy number per cell (*P* = 0.09, Mann–Whitney U test).

### Supplementary Tables

**Table S1: Susceptibility of strains used in the study to AZT**, expressed as minimum inhibitory concentration (MIC)

| **Mating Partner** | **Strain** | **MIC (µg/mL)** |
|---|---|---|
| **Donor** | AZT-susceptible *E. coli* MG1655::*mCherry* | 8 |
| | AZT-resistant *E. coli* MG1655::*mCherry* | 125 |
| **Recipient** | AZT-susceptible *E. coli* MG16555::*aacC1* | 8 |
| | AZT-resistant *E. coli* MG1655::*aacC1* | 125 |

**Table S2: Major single-nucleotide polymorphisms (SNPs) detected in the AZT-resistant donor and recipient strains** by comparison with susceptible strains

| **AZT- Resistant *E.coli* Strain** | **Mating Partner** | **Gene** | **Nucleotide Change** | **CDS pos. / CDS length** | **Amino acid change** |
|---|---|---|---|---|---|
| **MG1655::*mCherry*** | Donor | *yjjG* | G → A | 629/678 | Gln27* |
| **MG1665::*aacC1*** | Recipient | *tdk* | C → T | 79/618 | Trp210* |
